# Synthetic mRNA expressed Cas13a mitigates RNA virus infections

**DOI:** 10.1101/370460

**Authors:** Swapnil S. Bawage, Pooja M. Tiwari, Philip J. Santangelo

**Affiliations:** Wallace H. Coulter Department of Biomedical Engineering, Georgia Institute of Technology and Emory University, 950 Atlantic Dr. NW, Engineered Biosystems Building, Atlanta, GA, 30332, USA

## Abstract

The emergence of the CRISPR-Cas system as a technology has transformed our ability to modify nucleic acids. Prokaryotes evolved one member of this family, CRISPR-Cas effector, Cas13a, as an RNA-guided ribonuclease that protects them from invading bacteriophages. Here, we demonstrate that Cas13a can be programmed to target eukaryotic viral pathogens, influenza virus A (IVA) and human respiratory syncytial virus (hRSV) in human cells. We designed synthetic mRNA coding for Cas13a, which when guided by CRISPR RNAs (crRNA) to target influenza virus or hRSV RNA, significantly mitigates these infections both prophylactically, therapeutically, and over time. These data demonstrate a possible new class of synthetic mRNA-powered anti-viral interventions.

**One Sentence Summary:** crRNA guided Cas13a halts RNA virus infections

---

Recent outbreaks of Nipah (*1*), Zika (*2*) and Ebola (*3*), and the potential for future influenza pandemics (*4*), warrant the development of new classes of anti-viral drugs (*5*). Current drug development is focused on small molecules and neutralizing antibodies, which require high doses or frequent re-dosing to obtain functional outcomes (*6, 7*). Here, we propose a new paradigm for treating viral infections - the use of viral RNA activated RNases. RNA targeting enzymes can be advantageous over epitope blocking molecules. The stoichiometric limitations of antibody or small molecule neutralization can be overcome by utilizing RNA-targeting RNases, allowing them to “keep up” with rapidly proliferating viral systems. Here, we designed synthetic mRNA encoding Cas13a, guided by CRISPR RNAs (crRNAs) to target both genomic and messenger RNA of influenza virus A/WSN/1933(H1N1), and hRSV A2. Overall, we demonstrate mRNA expressed Cas13a and crRNA mediated mitigation of influenza and hRSV infections, both as prophylaxis and therapy.

The discovery of RNA targeting class II - type VI CRISPR-Cas system in bacteria has engendered tremendous interest for potential applications. The CRISPR locus is composed of an array of direct repeats (DR), spacer sequences and the gene encoding Cas protein (*8*). The effector type VI protein, Cas13a, processes DR of the precursor-crRNA (CRISPR-RNA) into mature crRNA. The Cas13a: crRNA complex activates when the target RNA (trRNA) compliments with crRNA, and the Cas protein initiates RNA cleavage due to the higher eukaryote and prokaryote nucleotide-binding (HEPN) domain (*8*). This property has been used to detect specific transcripts in mixtures of nucleic acids (*9*). Later, these findings were applied to develop a platform for viral and bacterial pathogen detection system, termed as High-Sensitivity Enzymatic Reporter UnLOCKing (SHERLOCK) system (*10, 11*). Cas13a activity has also been used for RNA knockdown of endogenous genes in human cell lines, and in the *Oryza sativa* (rice) protoplasm by transfecting with DNA vectors expressing Cas13a and crRNA (*12*). In addition, it was demonstrated that if *Nicotiana benthamiana* is infiltrated with Cas13a DNA, it can interfere with green fluorescent protein expressing Turnip mosaic virus (*13*). However, the utilization of this RNA targeting enzyme inhibiting RNA virus infections in human cells has not been reported.

For the RNA targeting RNases to be safe and effective for therapeutic use, rapid but transient expression is preferred (*14*). To achieve this, we opted for synthetic mRNA to express Cas13a. mRNA has the advantage of rapid translation of the desired protein and clearance, while it avoids safety concerns such as, genome integration, vertical and horizontal transmission, and long-term persistence in the body (*15*). We synthesized mRNA coding for *Leptotrichia buccalis* Cas13a (*9*) and designed multiple crRNAs (CR) and target RNAs (trRNA or TR) corresponding to influenza virus A WSN/33 (IVA) or hRSV A2 genome and mRNA. The crRNA consists of conserved DRs that are specifically recognized by *L. buccalis* Cas13a followed by an influenza or hRSV virus targeting sequence. The mRNA encoding Cas13a (modified with a V5 epitope tag) and Cas13a-NLS (with a V5 tag, and nuclear localization sequence each on C, and N termini) were modified during the *in vitro* transcription process to increase the translational efficiency and assuage innate immune responses.

Using the rabbit reticulocyte lysate, Cas13a and Cas13a-NLS mRNAs were translated *in vitro* and used to assess the RNA cleavage activity of Cas13a protein in conjunction with IVA crRNAs and trRNAs (Fig. 1A). crRNA and trRNA were derived from genome segments of IVA (Table S1). RNaseAlert™ substrate fluorescence was the output of RNA cleavage. Cas13a and Cas13a-NLS RNA cleavage generated fluorescence increased to its maximum during the initial 10 and 20 min period, respectively, and then gradually decreased over time due to depleted target RNA. The overall trend of RNA cleavage was similar for both the Cas13a (Fig. 1B) and Cas13a-NLS (Fig. 1C). RNA cleavage was also observed when the lysate was interrogated by gel electrophoresis using a 15% TBE-Urea gel (Fig. 1D and 1E). Our results indicate that *in vitro* translated Cas13a mediated RNA cleavage is specific and occurred only when both the crRNA and trRNA were present. Our results corroborate previous findings of specific activation, when purified Cas13a protein was used (*9*). The presence of yeast tRNA or cellular RNA in the cleavage buffer to assay Cas13a activity did not yield fluorescence or cleaved RNA products, demonstrating the specificity of Cas13a: crRNA towards target RNA. We observed that these designed crRNAs showed RNA cleavage with *in vitro* translated Cas13a and Cas13a-NLS only in the presence of corresponding trRNAs (figs. S1, S2).

**Fig. 1.**
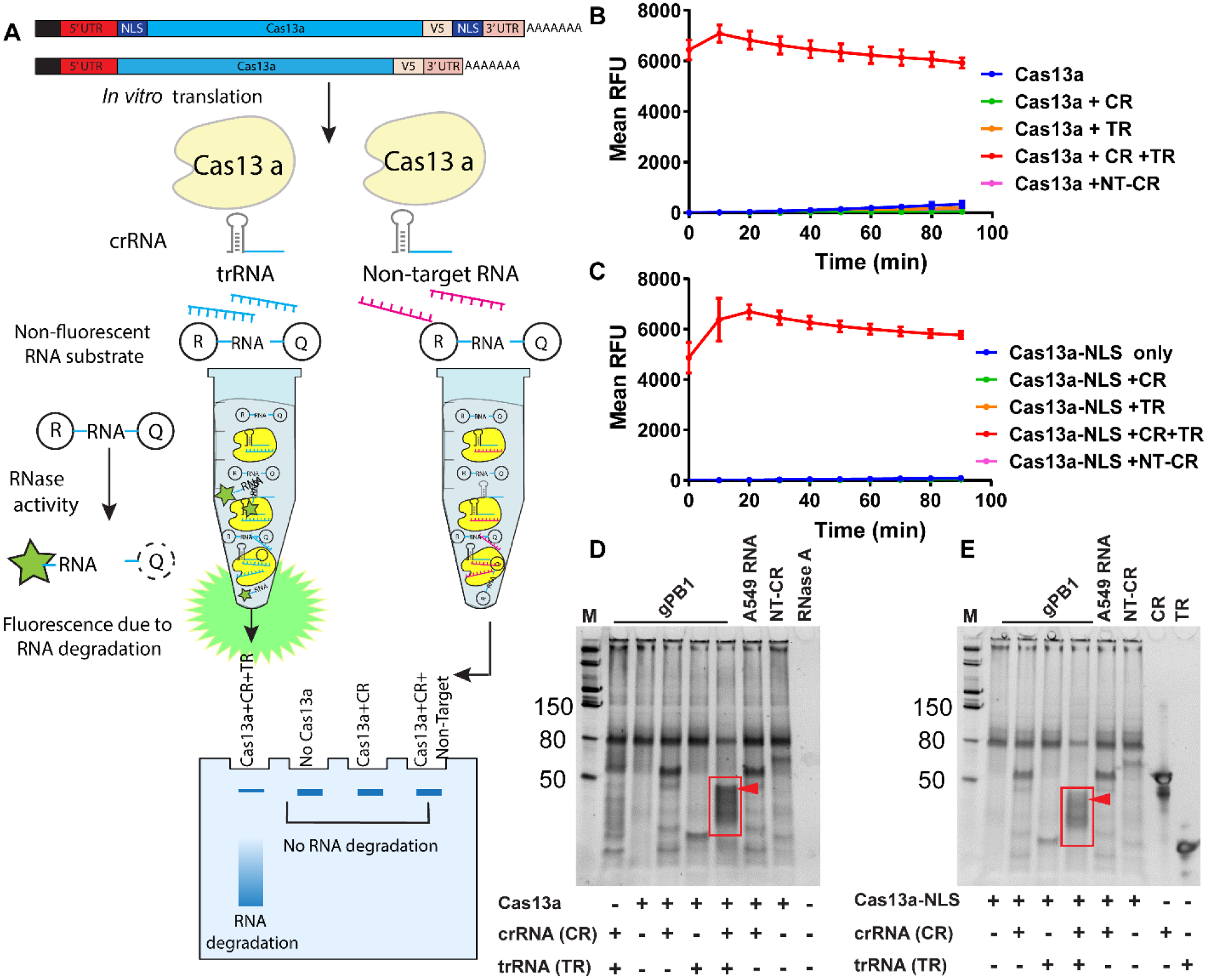
mRNA expressed *in vitro* translated Lbu Cas13a and Cas13a-NLS mediates target specific RNA cleavage. (A) Schematic representation of RNA cleavage assay using *in vitro* translation of Cas13a and Cas13a-NLS. *In vitro* transcribed mRNA expressing either Cas13a or Cas13a-NLS was translated *in vitro* using rabbit reticulocyte translation system. *In vitro* translated product was mixed with crRNAs, their corresponding target RNAs and an RNase alert substrate, which fluoresces upon cleavage, owing to RNase activity. (B) Target specific RNA cleavage of Cas13a and (C) Cas13a-NLS using crRNA (CR) targeting PB1 gene of influenza virus (gPB1) and its specific target RNA (TR). A non-target crRNA (NT-CR) was used a negative control. The RNA cleavage activity of Cas13a was measured at room temperature, for 90 minutes using mock translated product as a blank control. Mean RFU is represented as a means of triplicates. (D) A denaturing gel demonstrating RNA cleavage activity of Cas13a and (E) Cas13a-NLS against PB1 gene in the presence or absence of the corresponding CR and TRs.

These findings prompted us to assess the Cas13a:crRNA system in cellular models for influenza infections (Fig. 2A). First, we assessed mRNA delivery using transfection agents, Viromer^®^ Red, and Lipofectamine 3000, and via electroporation using the Neon^®^ transfection system in MDCK and A549 cell lines, all permissive to both influenza and hRSV. Cas13a expression was evaluated at 2, 4, 6, 16 and 24 h time points (fig. S3). Cas13a expression was observed as early as 2 h post transfection for all transfection agents and cell types. However, 16 h post transfection, the Cas13a expression decreased in the all cell lines transfected with Neon and Lipofectamine 3000. The Viromer^®^ Red transfected cells showed expression even at 24 h. Based on the results, we selected Viromer^®^ Red and A549 cell lines for all our subsequent experiments. Using immunofluorescence, we observed that Cas13a localized predominately within the cytoplasm, whereas Cas13a-NLS, was present in both the cytoplasm (as Cas13a is being synthesized) and, at 48 hr, within the nucleus (Fig. 2B).

**Fig. 2.**
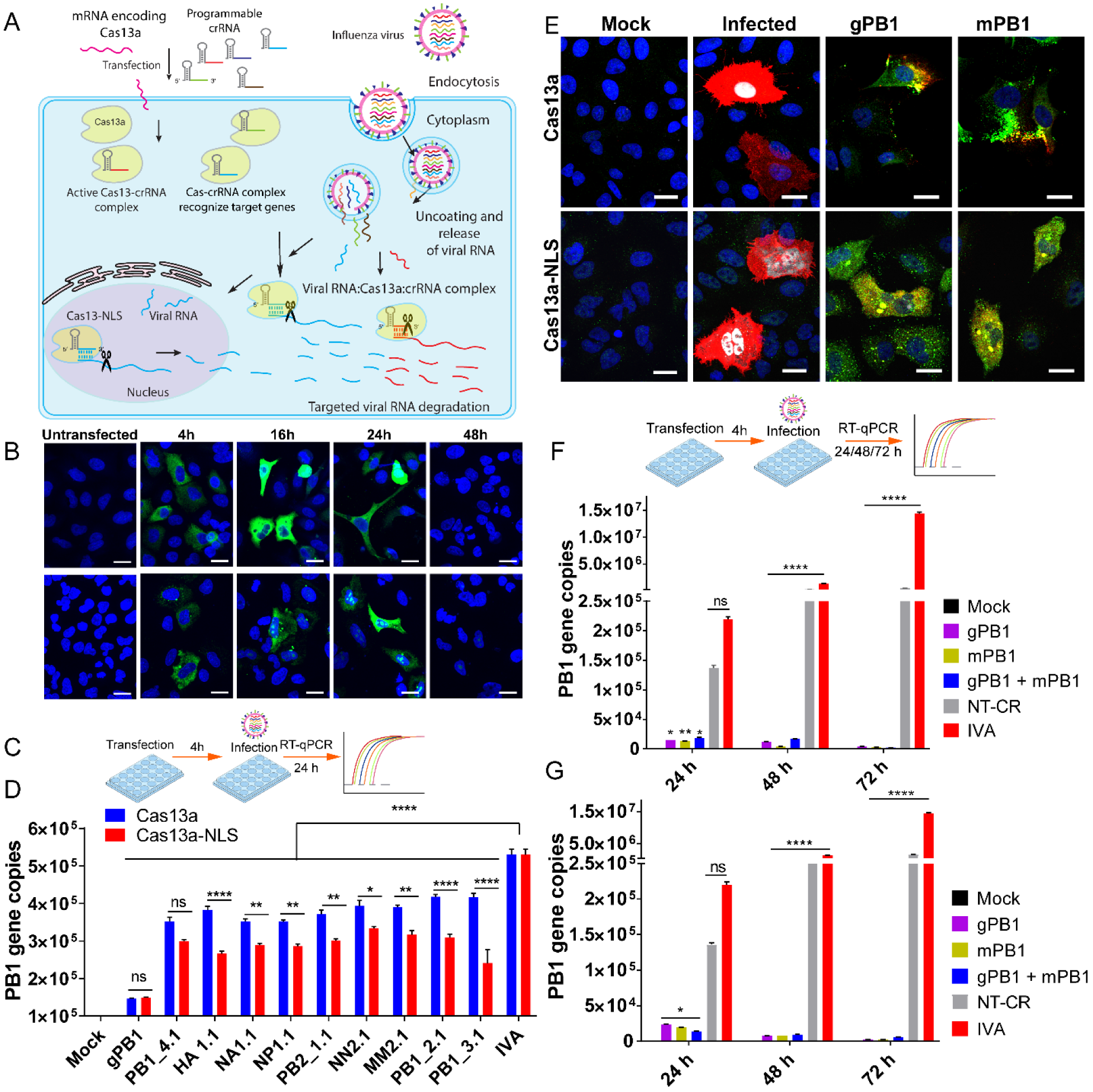
Cas13a and Cas13a-NLS express and target IVA specifically in cells. (A) Schematic representation of the mechanism of IVA RNA cleavage by Cas13a and/or Cas13a-NLS. (B)Expression kinetics of Cas13 and Cas13a-NLS in A549 cells using Viromer Red. Cas13 is stained using the V5 tag (green), DAPI is stained in blue. (C) Schematic of experimental set-up to test IVA inhibition in A549 cells in (D-G).) (D)RNA cleavage activity of Cas13a (blue bars) and Cas13a-NLS (red bars) with various crRNAs targeting IVA genes *in vitro* (A549 cells). (E) Cas13a and Cas13a-NLS inhibit IVA infection in cells transfected with mRNA encoding either proteins and crRNA for IVA gene PB1 genomic (gPB1) and messenger (mPB1) RNA, for 4h before infection. Cells transfected and infected were stained for IVA M2 protein (red), FISH targeting IVA (greyscale), Cas13a V5 (green) and nuclei (DAPI, blue). (F) Cas13 system prevents IVA infection up to 72 hours when transfected with either Cas13a or (G) Cas13a-NLS, and crRNAs targeting IVA PB1 gene (gPB1 and mPB1). Note the Cas13a system does not inhibit IVA infection when an unrelated crRNA is used as a control.

We then screened an array of crRNAs targeting multiple RNA segments to find candidate crRNAs that target and decrease IVA infections. A549 cells were simultaneously transfected with Cas13a or Cas13a-NLS mRNA and each crRNA followed by influenza virus infection (at multiplicity of infection, MOI 0.01) (Fig. 2C). Among all the crRNAs screened, the crRNA targeting the PB1 genome segment (gPB1) was found to reduce viral RNA copy most efficiently (Fig. 2D). To demonstrate that the Cas13a:crRNA system impacts viral RNA and protein production, we performed immunofluorescence for Cas13a, IVA M2 protein (indicating viral assembly or disassembly sites(*16*)), followed by fluorescence in situ hybridization (FISH) for the IVA genome. We observed that cells transfected with either the Cas13a or Cas13a-NLS mRNAs, and crRNA, targeting PB1 site, (gPB1 and mPB1 targeting PB1 mRNA) showed reduced IVA infection (Fig. 2E), and localization of Cas13a to viral sites within the cytosol. We next investigated the time dependent degradation of viral RNA during the infection. We found that targeting either the gPB1 or mPB1 reduced viral copies by 0.96-1.21 log at 24h, 1.88-2.6 log at 48 h, but further reduced viral RNA copies by 3.53-3.8 logs at 72h for both Cas13a (Fig. 2F) and Cas13a-NLS (Fig. 2G). This data is noteworthy, as prolonged effect of Cas13a interrupts IVA replication and reduced the viral copies up to 72h.

The Cas13a activity was specific in cleaving IVA RNA and did not result in any detected off-target RNA cleavage. The transcriptome profile of cells transfected with Cas13a mRNA and crRNA (PB1) at 8 and 24h post-delivery, with or without infection with IVA, showed no significant changes in endogenous gene expression (fig. S4). Similarly, specific endogenous mRNA knockdown in cells with no off-target activity was reported previously in *Leptotrichia wadei* Cas13a (*12*).

Next, we investigated a number of experimental conditions, in order to characterize the performance of the Cas13a:crRNA system. We evaluated the effect of MOI, demonstrating that Cas13a:crRNAs targeting PB1 were able to reduce the influenza virus gene copies by 0.77-1.26 log 24h post transfection(Fig. 3A), at MOI of 0.1 (Fig. 3B) and 0.01 (Fig. 3C), both physiologically relevant MOIs. We also observed that Cas13a: crRNA system effectively inhibits IVA in a normal human primary bronchial/tracheal epithelial cell model (Fig.3D). As we have demonstrated that the Cas13a: crRNA system mitigates IVA infections in a prophylactic manner, we sought to assess if the infection could be reduced in cells already infected with IVA. The cells infected with IVA (MOI 0.01) for 4h were later transfected with Cas13a/Cas13a-NLS mRNA and crRNA (gPB1 and mPB1) (Fig. 3E), and the effect on IVA monitored for 72 hours. We observed a maximum of 0.74 log reduction in IVA copy numbers with either Cas13a (Fig. 3F) or Cas13a-NLS (Fig. 3G) after 48 and 1.47 log reduction after 72h of transfection.

**Fig. 3.**
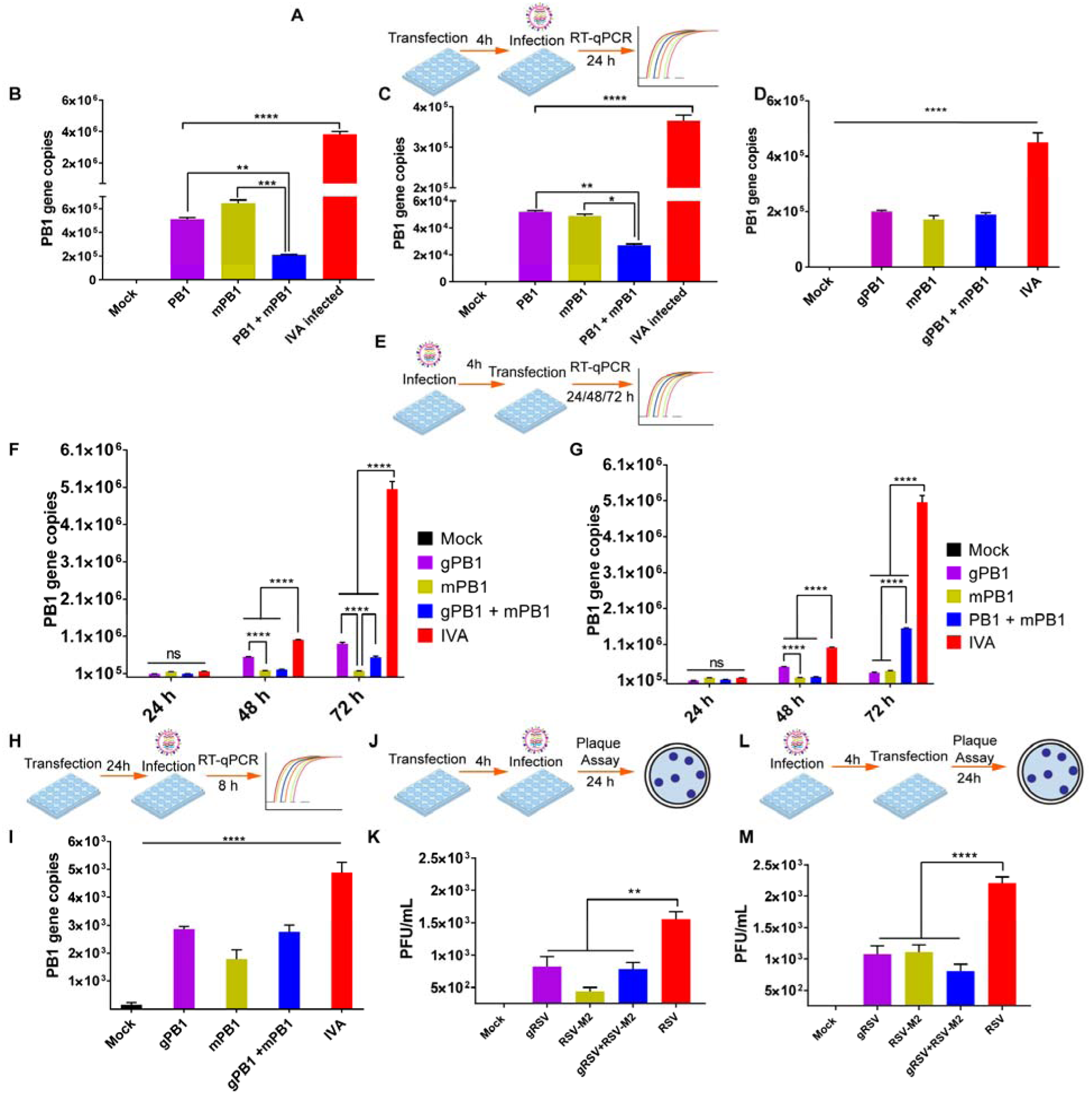
Cas13a is effective in preventing viral infection in different experimental set-up and can be adapted for other viruses e.g. RSV. (A) Schematic representation of experimental set-up. (B) Efficacy of Cas13a was tested at different MOI prophylactically at a multiplicity of infection of 0.1, and (C) at 0.01. (D) Cas13a also prevents IVA infection in human broncho-tracheal primary epithelial cells prophylactically. (E) Experimental set-up of infection before transfection with Cas13a. (F) In the infection first set-up. IVA infection was significantly reduced in both Cas13a, and (G) Cas13a-NLS system, for up to 72 h. (H) Schematic of experimental set up of delayed infection assay. (I) Cas13a inhibits IVA infection in a delayed infection model. Cells were transfected with Cas13 and CR gPB1 and mPB1 for 24 h and then infected with IVA for 8h. IVA gene PB1 copy numbers were significantly reduced in all treatment groups (I). The present Cas13a system was then adapted to prevent RSV infection, in both 4h transfection first-set-up (J-K) as well as 4h-infection first-set-up (L-M)). RSV infection was reduced up to 1-log when measured by plaque assay using Cas13a both prophylactically (K) and post-infection (M). Interestingly, RSV plaques in all treatment groups were reduced in both numbers and size. All the statistical analyses were performed using analysis of variance with multiple comparisons with appropriate post-test measurements. (* p≤ 0.05, **p≤ 0.005, ***p≤ 0.001 or ****p≤ 0.0001).

We then evaluated the efficiency of Cas13a-crRNA on the IVA infection by transfecting the cells with Cas13a mRNA and crRNA for 24h, anticipating a decrease in expression of Cas13a (based on Fig. 2B), and then infecting with influenza virus for 8h (Fig. 3H). Under this condition, we still observed a maximum of 0.3 log reduction of IVA, thus underlining the robustness of the Cas13a mediated targeting of IVA (Fig.3I).

Cas13a-crRNA system offers potential resolve to target various viral species through the design of specific crRNAs. To demonstrate this, we designed crRNA against both the human respiratory syncytial virus (RSV) genome (gRSV) and the M2 mRNA (RSV-M2). Cells transfected with Cas13a mRNA and both crRNAs exhibited reduced RSV titers, both as prophylaxis (Fig. 3J-K) and post-infection (Fig. 3L-M). There was maximum of 0.55 log reduction in Cas13a: crRNA when given prophylactically (Fig. 3K). However, when infected with RSV before transfection, Cas13a: crRNA reduced RSV titer by 0.3-0.44 log in crRNA 1.1 and m1.1, given together, whereas viral titer in crRNAs targeting genome mRNA (1.1) and RSV mRNA given individually was only 0.2-logs (Fig. 3M). It should be noted that even though we only observed ~50% knockdown via prophylaxis in hRSV, these data were obtained with a first-generation guide, based only on the M2/L gene end sequence. In addition, in the cells receiving Cas13a:crRNA, the plaques were not only diminished in number but also in size, demonstrating lower cell-to-cell spread and possibly the generation of more defective interfering particles. Overall, this result alone is extremely promising, as future unbiased guide screens against the hRSV genomic and mRNA will likely yield improved knockdown. These findings demonstrate the potential of the Cas13a: crRNA system as an antiviral approach, likely applicable to many viral pathogens.

Overall, our study clearly demonstrates that Cas13a mitigates IVA and hRSV infections and has the potential to be used against other RNA viruses. With the help of epidemiological information through screening and sequencing of the viral pathogens, several crRNAs can be rapidly and easily designed and screened to offer flexibility to deal with genetic drift and shifts or even the emergence of a new viral threat. However, it should be noted that the mRNP secondary structure and protein content, as well as viral RNP genome organization may influence the accessibility of crRNA and Cas13a for the target RNA. In our experiments, Cas13a cleavage of viral RNA was always successful, but viral RNPs, in the cell, were not always accessible. Thus, it is important to screen crRNAs in cells, considering these complexities; the development of high throughput screening methods for mining efficient crRNAs and those that address safety concerns will be critical to transforming this tool into a bona fide therapeutic approach.

## Acknowledgments

The authors acknowledge the help from Shweta Biliya and Fredrick Vannberg, Biomolecular Analysis Core, Georgia Institute of Technology for library preparation. We thank Anton Bryksin and Naima Djeddar, Molecular Evolution Core, Georgia Institute of Technology for RNA sequencing.

## Data and materials availability

All data is available in the main text or the supplementary materials. RNA-Seq data is in process of being deposited in SRA, NCBI.

## Supplementary Materials

Materials and Methods

Figures S1-S4

Tables S1-S3

External Databases S1

References (*9, 17*)

